# Augmenting the human interactome for disease prediction through gene networks inferred from human cell atlas

**DOI:** 10.1101/2024.12.12.628105

**Authors:** Euijeong Sung, Junha Cha, Seungbyn Baek, Insuk Lee

**Affiliations:** Department of Biotechnology, College of Life Science and Biotechnology, Yonsei University, Seoul 03722, Republic of Korea

**Author notes:** Corresponding author: Insuk Lee. These authors contributed equally to this work (Euijeong Sung, Junha Cha).

**Keywords:** Co-expression network, cell-type-specific gene network, single-cell atlas, the human interactome, disease gene prediction

## Abstract

Gene co-expression network inference from bulk tissue samples often misses cell-type-specific interactions, which can be detected through single-cell gene expression data. However, the noise and sparsity of single-cell data challenge the inference of these networks. We developed scNET, a framework for integrative cell-type-specific co-expression network inference from single-cell transcriptome data, demonstrating its utility in augmenting the human interactome for more accurate disease gene prediction. We address the limitations in *de novo* network inference from single-cell expression data through dropout imputation, metacells formation, and data transformation. Employing this data preprocessing pipeline, we inferred cell-type-specific co-expression links from single-cell atlas data, covering various cell types and tissues, and integrated over 850K of these inferred links into a preexisting human interactome, HumanNet, resulting in HumanNet-plus. This integration notably enhanced accuracy of network-based disease gene prediction. These findings suggest that with proper data preprocessing, network inference from single-cell gene expression data can be highly effective, potentially enriching the human interactome and advancing the field of network medicine.

## Introduction

The human body consists of various tissues, each organized by distinct cell types. Genes exhibit different expression programs and interact with different molecules depending on the tissue and cell type. Understanding these gene interactions is crucial for comprehending various mechanisms within the human body and gaining insights into the development of diseases that arise when these interactions are disrupted. Gene network models have been successfully applied to understand gene functions and predict their involvement in human diseases [1,2]. Extensive efforts to integrate network links inferred from various evidence have enabled the construction of genome-scale human interactome, significantly contributing to our understanding of human diseases.

Among many evidence types supporting gene functional associations, co-expression is probably the most widely used one because of availability of an enormous amount of gene expression data to public. Co-expression networks are generally inferred from bulk tissue expression data. This analysis cannot infer cell-type-specific co-expression network which can be detected from statistical test for expression variability between cells. Moreover, a recent study demonstrated that much of the co-expression based on bulk tissue expression data is attributable to the cellular compositional effects rather than intracellular regulatory relationship [3]. Thus, the human interactome may currently lack of cell-type-specific gene interactions. Addition of such interactions will enhance the human interactome in prediction of gene function and disease association through capturing cell-type-specific functional interactions. The single-cell RNA sequencing (scRNA-seq) technology enabled co-expression analysis between cells. By testing co-expression among cells of the same cell type, it allows for the inference of cell-type-specific co-expression links [4]. However, *de novo* network inference from scRNA-seq data is challenging due to its high-level of noise and sparsity [5,6]. Therefore, these inherent limitation of scRNA-seq data needs to be addressed to identify reliable co-expression links.

Here, we present scNET, a framework for integrative cell-type-specific co-expression network inference from single-cell gene expression data. We addressed the problem of noise and sparsity through of preprocessing of raw scRNA-seq data such as dropout imputation, metacell formation, and data transformation. The inferred co-expression links from the preprocessed single-cell transcriptome data are then evaluated and integrated into the preexisting human interactome. By applying scNET to various single-cell atlas data, we added over 850K cell-type-specific co-expression links to a preexisting human interactome, HumanNet [1], constructing “HumanNet-plus.” We demonstrated that HumanNet-plus significantly enhances the ability to predict disease-associated genes.

## Results

### scNET framework for de novo network inference from single-cell gene expression data

Inference of co-expression links from single-cell gene expression data suffer from their nature of high level of noise and sparsity. To overcome this hurdle, we capitalized three strategies of preprocessing single-cell transcriptome data: data transformation, dropout imputation, and metacell formation (**Fig. 1a**). Transformation of single-cell gene expression data by bigSCale (ver.2.0.0) [7,8] method allows detection of hidden correlations. Briefly, the bigSCale clusters cells with similar expression profiles then run differential expression analysis between all pairs of clusters. This process yields a Z-score for each gene. Co-expressions are detected through Pearson correlation coefficient (PCC) between these Z-scores. Imputation of dropouts also enhances detection of correlation between gene expression, although it also increases likelihood of presenting false links [9]. Thus, we assessed five dropout imputation methods for inference of gene functional associations : DCA [10], DeepImpute [11], scGNN [12], scIGAN [13], SAVER [14]. Pooling multiple cells makes analysis less susceptible to dropouts and noisy signals of single-cell data [15]. For this approach, we evaluated three algorithms of generating metacells: Metacell [16], Metacell2 [17], and SuperCell [18].

**Figure 1.**
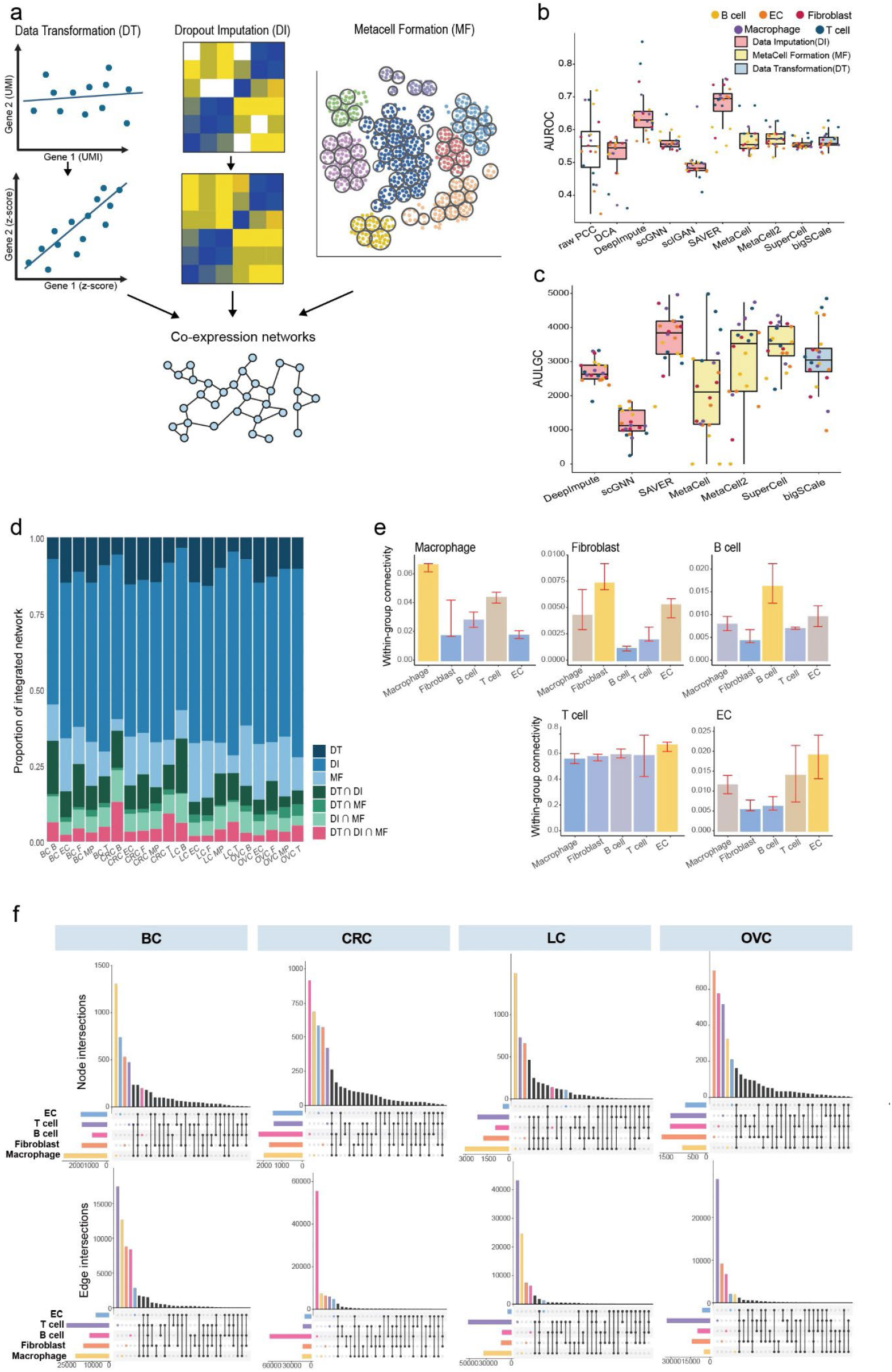
Overview of scNET framework. **a**. Schematic overview of *de novo* inference of cell-type-specific co-expression network through three strategies of preprocessing of single-cell gene expression data. **b-c**. Box plot of area under the receiver operating characteristic curve (AUROC) (b) and area under the log likelihood score against genome coverage (AULGC) (c). Different strategies of data preprocessing are marked by color bar. **d**. Stacked bar plot of three groups of co-expression links inferred from different data preprocessing strategies and their overlaps. **e**. Bar graph illustrating the within-group connectivity of each cell-type marker genes derived from CellMarkerDB within the corresponding cell-type-specific networks. Connectivity is normalized by total number of links of each network. Relevant cell type for the given network is marked by yellow bar. **f**. Upset plot depicting interactions of edges between cell-type-specific networks for each cancer type.

We compared these data preprocessing methods through network inference from pan-cancer single-cell atlas data that includes immune/stromal cells from breast cancer (BC), colorectal cancer (CRC), lung cancer (LC) and ovarian cancer (OVC) biopsies [19]. We annotated cells with five major cell types: macrophage, fibroblast, B cell, T cell, and endothelial cell (EC). For cells comprising each cell type, we measured PCC between genes based on expression matrix generated from all the data preprocessing methods. To select cell-type-specific co-expression links that are likely to support functional association between genes, we utilized log likelihood score (LLS) scheme which is based on Bayesian statistics framework [20] (**Methods**). By retaining only links with positive LLS values—indicating that the cell-type-specific co-expression links are likely functionally coupled—we finalized gene networks for each cell type across various cancer types.

We assessed networks inferred from pan-cancer single-cell atlas data, processed with various data preprocessing methods, using area under the receiver operating characteristics (AUROC) curve analysis (**Fig. 1b**). As expected, network inferred from raw single-cell expression data demonstrated poor performance in predicting gene functional associations. Except for DCA and scIGAN, which showed prediction capacity comparable to a random model, all other preprocessing methods were further evaluated using a precision-recall-like analysis, measuring the area under the LLS against genome coverage curve (AULGC) (**Fig. 1c**). Based on these assessment results, we selected bigSCale, SAVER, and SuperCell as a method of choice for each category of preprocessing approach. We found that the inferred co-expression networks by three categories of preprocessing methods are largely complementary (**Fig. 1d**), which suggests that integrating co-expression links from the three approaches will increase comprehensiveness of the network. Therefore, we generated a network for each cell-type by integrating links inferred from preprocessed data using bigSCale, SAVER, and SuperCell through a weighted-sum method we previously developed [20] (**Methods**).

We evaluated cell-type-specificity of networks inferred from pan-cancer single-cell atlas data using marker genes for each cell type derived from the CellMarker database [21]. Our analysis revealed that connectivity among marker genes within each cell type (within-group connectivity) is generally highest in the corresponding cell-type-specific gene network (CGNs) (**Fig. 1e)**. Additionally, marker genes for each cell type occupy the largest proportion of network nodes within their corresponding CGNs (**Supplementary Fig. 1**). These results suggest that the *de novo* networks inferred using scNET framework accurately represent cell-type specific biology. Furthermore, when comparing various CGNs within each cancer type, we observed the highest proportion of edges unique to a single cell type across all cancer types (**Fig. 1f**). This indicates that even within the same tumor tissue, genes exhibit distinct interactions, reflecting their cell-type-specific functions.

### A compendium of gene networks across multiple tissues and cell types

With scNET framework, we constructed a compendium of CGNs across multiple tissues and cell-types. We separately collected normal cross-tissue immune cells from Human Cell atlas [22-24], normal stromal/epithelial cells from the Tabula sapiens project [22,24], and normal middle temporal gyrus (MTG) cells from single nucleus RNA-seq data of the Allen Brain Atlas [24]. While the majority of scRNA-seq datasets yielded CGNs, only a few snRNA-seq datasets for brain cells produced CGNs. In total, we successfully generated 100 CGNs across various tissues and cell-types (**Supplementary Table 1**).

To examine the interrelationship among the compendium CGNs, we represented each network with binary gene profiles (indicating the presence or absence of genes) and visualized them in a reduced dimensional space using Uniform Manifold Approximation and Projection (UMAP). This analysis revealed a clustering pattern where networks associated with similar cell types tended to be closer to each other (**Fig. 2a**). These results suggest that the CGNs in the compendium accurately reflect the biological properties of the corresponding cell types. In addition, we found that CGNs related to the same tissue types did not exhibit a clustering trend (**Fig. 2b**), indicating that cell type identity, rather than the tissue environment, predominantly determines the structures of CGNs.

**Figure 2.**
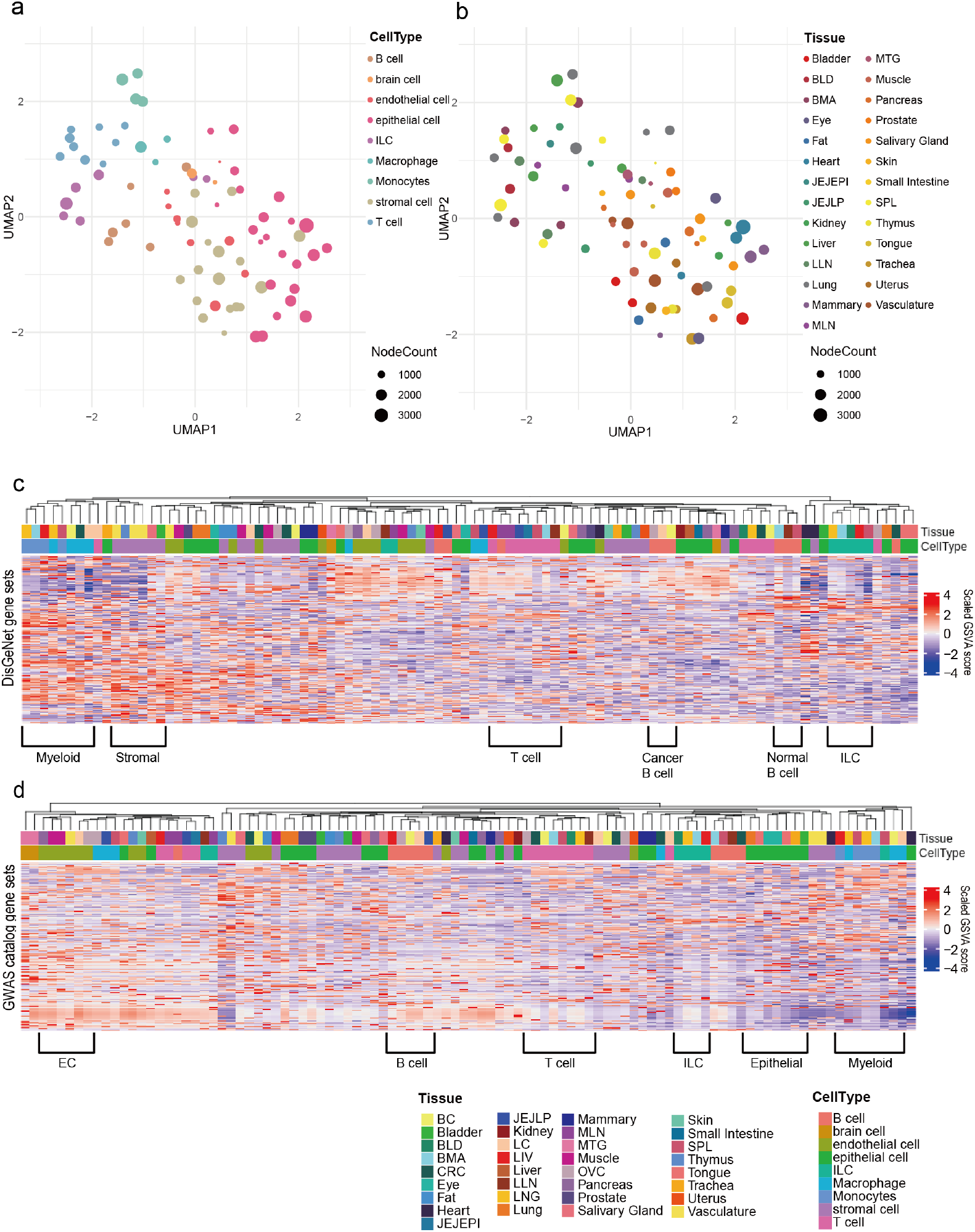
**a-b**. UMAP plots depicting the interrelationship among CGNs, with CGNs color-coded by cell types (a) or tissue types (b). **c-d**. Heatmaps showing the disease association of CGNs based on disease gene sets derived from the DisGeNET (c) and GWAS catalog (d) databases. Disease association was evaluated using GSVA scores. The scaled GSVA scores are indicated by color codes. CGNs were clustered based on the Spearman correlation of their disease-association profiles.

In many cases, gene expression influences human disease through specific cell types. Therefore, we expect that the cell-type-specificity of the compendium networks will positively contribute to the genetic dissection of human diseases. To evaluate this, we assessed how well the compendium GCNs reflects the cell type specificity of various diseases. Specifically, we investigated whether the compendium CGNs can deconvolute known disease genes to cell types using DisGeNET [25] and the GWAS Catalog [26]. Following the prioritization of genes within each CGN based on network degree centrality, we performed gene set variation analysis (GSVA) [27] to generate disease association score profiles, which were used to cluster CGNs. We observed that CGNs clustered according to the same or closely related cell types based on disease association profiles, regardless of tissue types, when analyzed by disease profiles based on DisGeNET (**Fig. 2c**) and the GWAS Catalog (**Fig. 2d**). These results suggest that scNET framework provides novel insights into cell-type-specificity across diseases by enabling the construction of reliable cell-type specific network models.

### Cell-type-specific links enhance the human interactome for disease prediction

Most human interactomes have been constructed without considering cell-type-specific contexts. As a result, functional interactions that occur exclusively within particular cell types have often been overlooked in conventional network inference approaches, such as experimental detection of protein interactions and co-expression analysis using bulk tissue samples. The scNET framework, which enables reliable *de novo* identification of cell-type-specific co-expression links, offers an opportunity to integrate these links into the preexisting human interactome, potentially enhancing its capability for disease prediction.

To test this, we integrated the compendium CGNs into a human interactome, HumanNet (version 3) [1], and evaluated the increase in disease predictivity of the integrated network compared to the original interactome. We first constructed a single cell-type-specific network by integrating the 100 CGNs in the compendium using a weighted sum scheme (**Methods**). This combined network was subsequently integrated with multiple component networks of HumanNet, resulting in an enhanced version that we refer to as HumanNet-plus. Finally, we selected 1,980,000 interactions that are more likely to occur than by random chance for HumanNet-plus, of which 854,506 links (43.2%) were derived from compendium CGNs.

Typically, temporally consistent and spatially prevalent gene-gene functional interactions have higher likelihood scores compared to dynamic and cell-type-specific ones. Therefore, the newly incorporated links from the CGNs are expected to exhibit relatively low likelihood scores, as they represent functional interactions occurring only in specific cell types. As anticipated, the newly incorporated network links in HumanNet-plus predominantly showed low likelihood scores (**Fig. 3a**).

**Figure 3.**
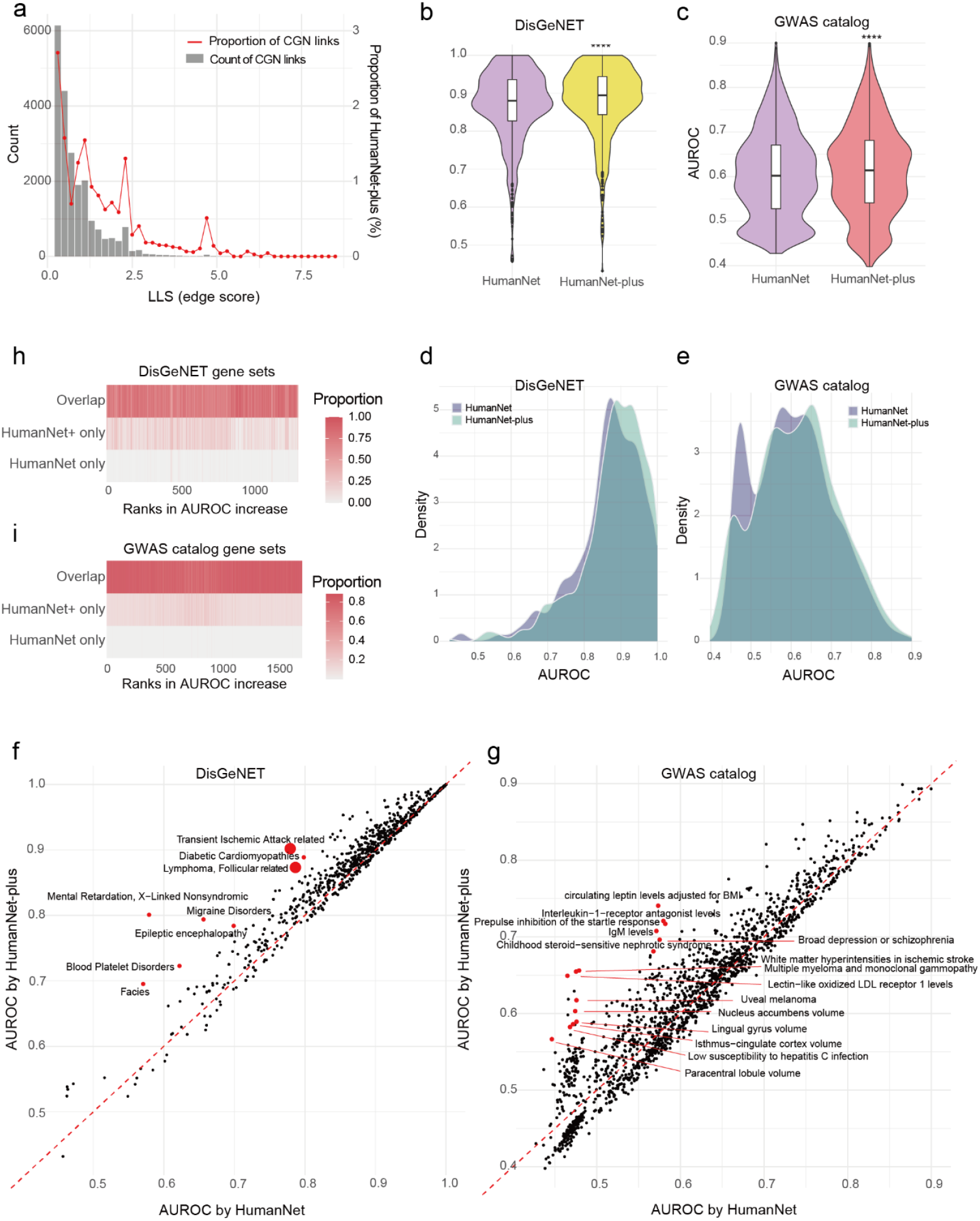
**a**. Distribution of edge scores (LLS) of CGN link of HumanNet-plus, presented in terms of count values and percentage proportions. **b-c**. Comparison of disease predictivity between HumanNet and HumanNet-plus using AUROC for disease gene sets derived from the DisGeNET (b) and GWAS catalog (c) databases. Significance was evaluated by two-tailed Mann-Whitney U test. (**** *P* < 0.0001) **d-e**. Density distribution plots comparing the AUROC for disease gene sets derived from the DisGeNET (d) and GWAS catalog (e) databases between HumanNet and HumanNet-plus. **f-g**. Scatter plots of AUROC scores comparing HumanNet (x-axis) and HumanNet-plus (y-axis) using DisGeNET (d) and GWAS catalog (e) disease gene sets. The top 15 diseases showing the largest increase in AUROC in HumanNet-plus compared to HumanNet are marked by red dots. Larger dots represent the overlap of multiple data points. **h-i**. Heatmaps showing the proportion of each category of links—overlap, HumanNet only, and HumanNet-plus only—within the group links of each disease according to the DisGeNET (h) and GWAS catalog (i) database. Disease gene sets are ranked by the increase in AUROC scores.

To evaluate the improvement of the human interactome through the incorporation of cell-type-specific co-expression links, we compared HumanNet and HumanNet-plus in disease prediction using disease gene sets compiled from DisGeNET and the GWAS catalog databases. The prediction accuracy for each disease was assessed using the AUROC for the retrieval of disease genes by guilt-by-association across disease gene sets containing at least 10 member genes. From these analyses, we observed a significantly higher AUROC for HumanNet-plus compared to HumanNet when using both disease gene databases (**Fig. 3b-c**). The AUROC scores were particularly increased for disease gene sets derived from the GWAS catalog (**Fig. 3d-e**).

We further examined individual diseases with improved predictions in HumanNet-plus and found that many of them are associated with disorders caused by the functional impairment of specific cell types (**Fig. 3f-g**). When assessing the overlap of network edges within disease genes across the networks, we found a substantial overlap between HumanNet and HumanNet-plus. Notably, the edges unique to HumanNet-plus were enriched in disease gene sets that showed a marked increase in AUROC compared to HumanNet, indicating a significant contribution of the cell-type-specific co-expression links to the improved disease predictions (**Fig. 3h-i**). These results collectively demonstrated that cell-type-specific co-expression links inferred through the scNET framework can effectively enhance the disease predictivity of preexisting human interactomes.

## Discussion

Considering that intercellular variation in gene expression within the same cell type can reveal cell-type-specific co-expression networks, the increasing availability of scRNA-seq data presents significant opportunities in network biology. Moreover, given the proven utility of the human interactome in studying human diseases, single-cell network inference could greatly advance network medicine. In this study, we developed scNET, a computational framework for inferring *de novo* gene functional interactions from co-expression across single cells. This framework addresses current challenges in single-cell network inference by overcoming data sparsity through three key approaches: data transformation, dropout imputation, and metacell formation. After benchmarking various methods in each category, we identified bigSCale, SAVER, and SuperCell as the optimal methods for the initial inference of co-expression links from single-cell transcriptome data. We found that networks inferred by these methods are largely complementary, producing a more comprehensive network through data integration using the LLS scheme leveraging Bayesian statistics. As a result, we established an effective and reliable bioinformatic pipeline for cell-type-specific network inference from single-cell gene expression data.

Previous studies evaluated various metrics for measuring the correlation of single-cell gene expression in identifying gene interactions [5,28]. Although the results varied, there was a consensus that no effective metric currently exists for measuring single-cell gene expression correlations. Even the best metrics achieved accuracy only marginally better than random expectation. Instead of focusing on correlation metrics, we prioritized data preprocessing to improve single-cell network inference. We found that with proper data preprocessing, widely used correlation measures can perform effectively. From these findings, we conclude that data preprocessing is more critical than the choice of correlation metrics for successful single-cell network inference.

Recently, many atlas-scale scRNA-seq datasets across various tissues and cell types have been made publicly available. We hypothesized that cell-type-specific networks inferred from these atlas datasets could enhance the existing human interactome. To test this, we applied the scNET framework to atlas datasets covering a wide range of cell and tissue types, both in normal and diseased states, generating a compendium of 100 CGNs. These networks were found to reflect the characteristics of their corresponding cell types, including disease associations. When we integrated the 100 CGNs into a preexisting human interactome, HumanNet, the network size increased substantially. This integrated network, termed HumanNet-plus, was evaluated for its disease predictivity and demonstrated a significantly improved ability to identify disease genes. Since most human interactomes lack cell-type-specific functional interactions, this approach is likely to enhance disease prediction in other human interactomes as well. Therefore, we anticipate that the scNET framework, combined with the growing repository of single-cell atlas data, could significantly advance network medicine in the future.

## Methods

### Single-cell gene expression data

We utilized scRNA-seq and snRNA-seq datasets for this study. First, we used pan-cancer scRNA-seq data generated from breast cancer, colorectal cancer, lung cancer, and ovarian cancer patients [29]. We conducted manual cell type annotation for this dataset. Next, we used immune cell atlas data cross 16 tissues from 12 deceased donors [23] and the Tabula Sapiens dataset [22]. We additionally used snRNA-seq data from brain cells of the Allen Brain Atlas [24]. We used pre-annotated cell type information for these immune and brain cell atlas datasets.

### Assessing scRNA-seq data preprocessing methods for co-expression network inference

We used three approaches of scRNA-seq data preprocessing for inferring co-expression gene network: dropout imputation, metacell, and data transformation. We tested the following dropout imputation methods: DCA [10], DeepImpute [11], scGNN [12], scIGAN [13], SAVER [14]. Before running imputation tools, we filtered out genes that have zero values for more than 95% of total cells. For gene-gene correlation analysis with imputed gene expression data by SAVER method, we used the function *cor*.*genes()* provided by the SAVER package. For metacell approach, we tested the following metacell generators: Metacell [16], Metacell2 [17], and SuperCell [18]. Similarly, we filter out genes that have zero value for more than 95% of the total metacells. For data transformation approach, we used bigSCale (ver.2.0.0) [7,8]. We used *compute*.*network()* function and calculated Z-scores of genes which are subsequently used for correlation analysis. Briefly, this method constructs a probabilistic model through comparing gene expressions between groups of cells that represents the likelihood of an expression change, yielding a Z-score of genes for every combination of comparison between cell groups. Correlations between these Z-scores represent a transformed association metric between genes within the single cell data. Co-expression was evaluated by Pearson correlation coefficients (PCC) using *cor()* function of R package. We measured likelihood of inferred functional associations between genes with the given evidence, degree of co-expression based on PCC, using Bayesian statistics framework we previously developed [20]. We calculated log likelihood score (LLS) for every bin of 1000 inferred gene-gene associations sorted from the highest PCC score using the following equation,

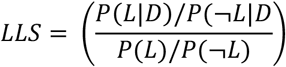

where P(*L*|*D*) and P(*¬L*|*D*) represent the probabilities of positive and negative gold-standard gene pairs for the given data, respectively, and P(*L*) and P(*¬L*) represent the probabilities of gold-standard positive and negative gene pairs, respectively. The gold-standard positive and negative gene-gene associations were derived from those used for modeling HumanNet [1]. Using regression model between PCC and LLS, we converted PCC into LLS for all inferred links. Finally, we selected gene-gene links with positive LLS which means the probability of functional association between these two genes is more likely than those by random chance.

We then assessed resultant networks for prediction of gold-standard gene-gene associations based on the receiver operating characteristics (ROC) analysis. We summarized performance by ROC analysis by area under the ROC curve (AUROC) which was analyzed by using R package ROCit. For precision-recall-like analysis, we used LLS and genome coverage which correspond to precision and recall, respectively. We summarized the performance by analysis using LLS and genome coverage using area under the LLS against genome coverage curve (AULGC).

### scNET: integrative cell-type-specific co-expression network inference framework

To establish an effective pipeline for identifying co-expression links from scRNA-seq data, we selected three data pre-processing methods, one from each category: bigSCale [7,8] (ver.2.0.0) for data transformation, SAVER [14] (ver.1.1.2) for data imputation, and SuperCell [18](ver.1.0.0) for metacell approach.

Briefly, bigSCale computes differential expression between minor clusters of cells grouped to reduce noise. Cells within each cluster are assumed to be biological replicates. With these N clusters × M genes matrix,

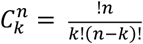

each pairwise computation between N clusters are performed to yield a Z-score matrix with 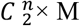 dimensions via *compute*.*network()*. ‘recursive’ was selected for parameter *clustering* and 0.9 for *quantile*.*p* parameter. The Z-score is then computed for PCC by R package function *cor()*.

For data imputation using SAVER, low expression values are inferred assuming gene expression profiles follow a negative binomial distribution with a Poisson-gamma mixture :

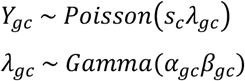

where *Y*_*gc*_ is the observed expression count (UMI) of gene g in cell c. and *λ*_*gc*_ represents the normalized “true” expression. After removing low expressed genes (over 95 % of cells with zero UMI count), SAVER imputed gene × cell matrix was calculated for gene-gene PCCs.

The supercell algorithm merge highly transcriptionally similar cells modifying the metacell algorithm. Briefly with cells as nodes and edges based on Euclidean distance, cells are grouped by walk_trap clustering algorithm. The graining level is consistently set to 20 across all datasets. Output matrix was then computed for gene-gene PCCs with *cor()*.

We next integrated three gene-gene correlation matrix using likelihood scoring and weighted sum method [20]. After inferring co-expression links through bigSCale, SAVER, and SuperCell methods, we convert the correlation score, PCC values, into LLS as described above. The three co-expression networks were integrated through WS score defined as the following equation:

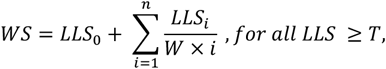

where *LLS*_0_ indicates the maximum LLS for the gene pairs and *LLS*_*i*_ are sorted LLS scores by decreasing order. The weight factor, W, and LLS threshold, T, are optimized for maximizing the area under the plot of LLS versus gene coverage. The WS score for each gene pair is then re-scored using the same LLS scheme.

### Disease profiling of CGNs

First, we selected shared genes among all 100 CGNs inferred from single-cell atlas data. We then ranked the shared network genes by degree centrality. Disease gene sets, comprising a total of 5,763 genes, were gathered from DisGeNET [25] and the GWAS Catalog [26] for analysis. The association of disease gene sets with each CGN was assessed using GSVA, performed through the *gsva()* function from the GSVA package [27]. Then we measure correlation of GSVA profiles of CGNs using Spearman Correlation Coefficient.

## Data availability

Network edge information of HumanNet-plus and compendium CGNs are downloadable from https://www.inetbio.org/humannet/humannet-plus/. All scRNA-seq datasets used in this study were downloaded from public resources. Raw sequencing data of the pan-cancer scRNA have been deposited in the ArrayExpress database at EMBL-EBI (www.ebi.ac.uk/arrayexpress) with accession numbers E-MTAB-8107, E-MTAB-6149 and E-MTAB-6653. Cross-tissue immune cell atlas data was downloaded from https://www.tissueimmunecellatlas.org/. Tabula Sapiens atlas data can be downloaded from GEO under accession number GSE201333. Brain snRNA-seq data was accessed at Allen Brain Map (https://portal.brain-map.org/atlases-and-data/rnaseq).

## Code availability

The R and Python codes used in this study are available on GitHub. (https://github.com/netbiolab/scNET)

## Acknowledgements

This research was supported by the Bio & Medical Technology Development Program of the National Research Foundation funded by the Ministry of Science and ICT (2022M3A9F3016364, 2022R1A2C1092062 to I.L.). The work was supported in part by Brain Korea 21(BK21) FOUR program.

## Author Contributions

E.S., J.C., and I.L. conceived and designed the study. J.C. built bioinformatic pipelines for the network inference and integration. E.S. performed data analysis and network construction. S.B. provided critical technical advice on the study. I.L. supervised research. E.S. and I.L. wrote the manuscript.

## Competing Interests

The authors declare no competing interests.

## Notes

### Competing Interest Statement

The authors have declared no competing interest.

