## supplementary figure 1 for "Augmenting the human interactome for disease prediction through gene networks inferred from human cell atlas"

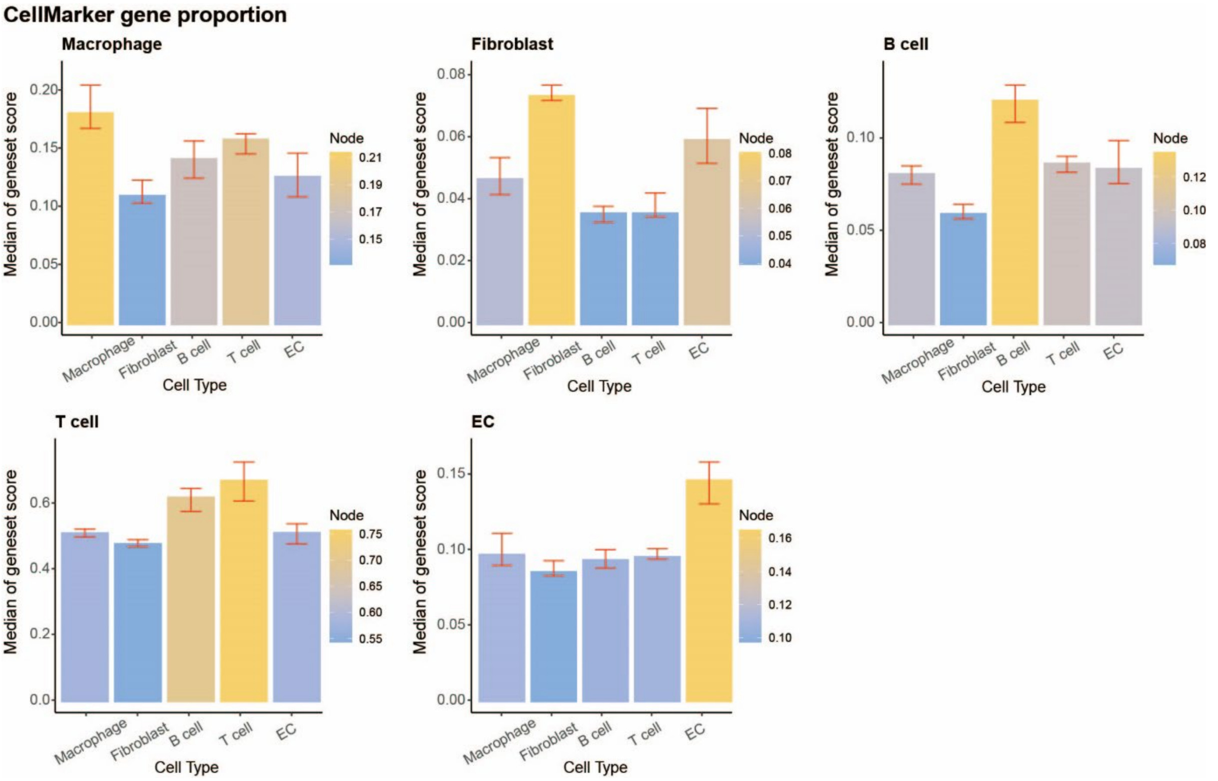


**Supplementary Figure 1**. Proportion of cell type-specific marker genes in the corresponding cell-type-specific network. The bar graphs represent the median proportion across four cancer types (breast, colorectal, lung, and ovarian cancers), with error bars indicating the range of scores across these cancer types.
